# *Trans*MPRA: A framework for assaying the role of many *trans*-acting factors at many enhancers

**DOI:** 10.1101/2020.09.30.321323

**Authors:** Diego Calderon, Andria Ellis, Riza M. Daza, Beth Martin, Jacob M. Tome, Wei Chen, Florence M. Chardon, Anh Leith, Choli Lee, Cole Trapnell, Jay Shendure

## Abstract

Gene regulation occurs through *trans*-acting factors (*e.g.* transcription factors) acting on *cis*-regulatory elements (*e.g.* enhancers). Massively parallel reporter assays (MPRAs) functionally survey large numbers of *cis*-regulatory elements for regulatory potential, but do not identify the *trans*-acting factors that mediate any observed effects. Here we describe *trans*MPRA — a reporter assay that efficiently combines multiplex CRISPR-mediated perturbation and MPRAs to identify *trans-*acting factors that modulate the regulatory activity of specific enhancers.

## Main

Cells rely on complex gene-regulatory networks in the context of differentiation, development, homeostasis, external signal response, etc^1–4^. These networks depend on myriad direct and indirect interactions between *trans*-acting factors and *cis*-regulatory elements, which underlie the recruitment of transcriptional machinery to proximally located genes. Across all genes, the fine-tuned orchestration of gene expression through such regulatory interactions enables an enormous diversity of cellular states^5,6^.

Despite the centrality of *trans*-acting factors to gene regulation, we lack robust methods for identifying <u>which</u> *trans-*acting factors mediate the functionality of <u>which</u> *cis*-acting regulatory elements. High-throughput methods such as MPRAs^7–10^ or CRISPR-QTL^11^ functionally validate putative enhancers or identify their target genes, but do not identify the *trans-*acting factors that mediate those effects. Gene perturbation screens^12,13^ identify *trans-*acting factors that directly or indirectly alter gene expression, but not the specific enhancers through which those effects are mediated. ChIP-seq^14^ and CUT&Tag^15^ profile the locations of a protein of interest genome-wide, but are biochemical rather than functional in nature. Targeted pulldown coupled to mass spectrometry can identify which proteins physically associate with a locus of interest, but such approaches do not readily scale^16–18^.

To address this gap, we developed the *trans* massively parallel reporter assay or *trans*MPRA. Here we describe *trans*MPRA together with a proof-of-concept in which we apply it to test all possible regulatory interactions between 8 *trans*-acting factors and 95 putative enhancers.

We first developed an iterative cloning strategy in which random combinations of guide RNAs (gRNAs; for CRISPR perturbation) and enhancers (for MPRA) are cloned to different parts of a bifunctional vector, but in such a way that the combination is compactly encoded in the functional readout of a STARR-seq-like^8^ MPRA (**Fig. 1a-c**; **Fig. S1**). In brief, a library of gRNA spacers and a library of barcodes are cloned adjacent to one another. PCR amplicons derived from this library are deeply sequenced in order to associate gRNAs with the specific barcode sequence(s) to which they are paired in the library. After introducing a constant sequence corresponding to a minimal promoter^19^, a library of enhancers is cloned to a site adjacent to the barcode. The resulting library is bifunctional, with each construct encoding both a Pol3-driven gRNA as well as an enhancer with the potential to drive its own transcription from an adjacent Pol2-driven minimal promoter. A key aspect of this MPRA design is that resulting mRNAs encode the identity of the enhancer (its own sequence, like STARR-seq^8^), as well as the sgRNA to which it is linked (the barcode).

**Fig. 1:**
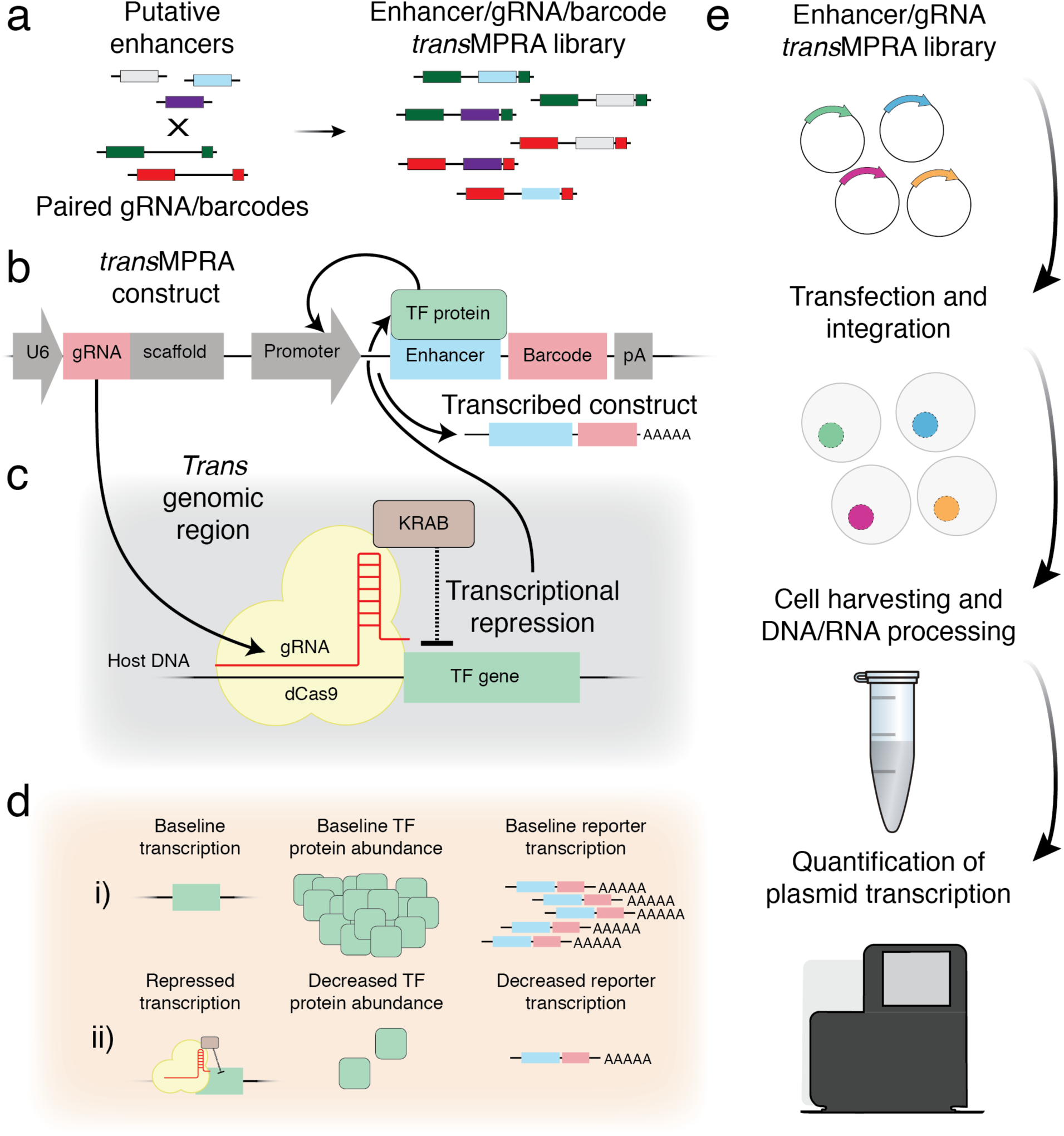
Overview of *trans*MPRA. **a**, Putative enhancers are cloned between gRNAs and gRNA-linked barcodes, resulting in a combinatorial library of enhancer/gRNA combinations. **b**, A representative *trans*MPRA reporter construct (pA, polyadenylation site). Critically, the resulting mRNAs encode the identity of both the enhancer (its own sequence) and the sgRNA to which it is linked (the barcode). **c**, Expressed gRNA directs the dCas9-KRAB complex to repress activity of the target TF. **d**, A sequencing-based readout differentiates two possible outcomes of any given knockdown-enhancer pairing. **e**, Schematic of the *trans*MPRA experimental workflow.

Rather than relying on transient transfection as is typical for MPRAs, we integrate the *trans*MPRA library into a dCas9-KRAB-expressing cell line using piggyBac transposase^20^. Integration allows the dCas9-KRAB complex sufficient time to reduce the transcript and protein levels of its targets^21,22^. In addition, it avoids the template switching associated with lentivirus, which would scramble the associations between gRNAs and their barcodes^23^.

Once the construct is integrated and the gRNA expressed, we hypothesize two possible scenarios (**Fig. 1d**). We assume an unknown set of protein factors underlie the ability of an enhancer to regulate gene expression. If the gRNA targets a protein that does not play a role in mediating the activity of the enhancer to which it is linked, then we expect no change to the enhancer-associated reporter activity. Alternatively, if the gRNA targets a protein that does play such a role, then we expect differential transcription of the enhancer’s reporter.

As is typical in MPRAs and to account for knockdown effects on cell proliferation, we sequence the self-transcribed enhancer element, together with the barcode that uniquely identifies the upstream gRNA, separately from both DNA and RNA (**Fig. 1e**). We then use the resulting counts to estimate the differential activity of each enhancer in the context of each encoded CRISPRi-mediated knockdown.

As a proof of concept, we designed a *trans*MPRA experiment to measure potential interactions between 8 *trans*-acting factors and 95 putative enhancers. Altogether, the enhancer library consisted of 101 regions, each 201 bp in length: 75 putative enhancers with high activity (‘positive regulators’) and 20 regions associated with low or no activity (‘weak regulators’) in K562 cells, as determined by a previous MPRA study^24^, and 6 scrambled versions of positive or weak regulators (3 of each; ‘scramble’) (**Table S1**).

We also identified 8 transcription factors (TFs) that were both expressed in K562 cells^25^ and had at least one significant motif match in one or more of the putative enhancers. These were *ATF4, FOSL1, GABPA, GATA1, MYC, NRF1, SP1*, and *STAT1*. We then selected 3 gRNAs to target the promoter of each of these 8 TFs via CRISPRi^26^, as well as 3 scrambled no-target gRNAs. One gRNA that targets *NRF1* was excluded prior to cloning because it contained a necessary restriction enzyme cut site, such that there were 26 gRNAs in total.

We next applied the aforedescribed iterative cloning strategy to combinatorially pair these gRNAs and enhancer fragments (26 x 101 = 2,626 possible pairings), while also introducing a degenerate 18 bp barcode (**Fig. S1**). During the association step, we identified 1.8 million unique barcodes (mean ∼68,000 per sgRNA; **Fig. S2**), indicating that the library construction strategy is capable of achieving high complexity.

A plasmid encoding the piggyBac transposase was transfected along with our plasmid library into three replicate samples of ten million K562 cells that constitutively express the dCas9-KRAB complex. We performed a GFP-based optimization experiment (**Fig. S3**), which led us to choose two library concentrations to test in parallel: 1) A higher multiplicity-of-integration (MOI) condition that resulted in an average of two integrations per cell, and; 2) a lower MOI condition at 20% of the higher MOI plasmid concentration (**Fig. S4**). We harvested aliquots of five million cells on day five (D5) and day ten (D10) post-transfection, extracting both DNA and RNA from a total of 12 samples (2 conditions x 2 timepoints x 3 replicates).

Each library was processed with a two-step PCR amplification strategy which introduced a library-specific sequencing index, a unique molecular identifier (UMI) and P5/P7 flow cell adapters (**Fig. S5**). Amplicons were pooled, size-selected, and deeply sequenced. We obtained 170 million reads passing QC and aligning to the *trans*MPRA construct. On average, each DNA library had 3.8 million reads and each RNA library had 10.2 million reads. Individual enhancer fragments, gRNAs, and enhancer/gRNA pairs were well represented (**Fig. S6**).

As to our knowledge, MPRAs have not previously been conducted via piggyBac integration, we first sought to validate that the MPRA was successful by focusing on the subset of data from reporters bearing a scrambled control gRNA (**Fig. 2**). To estimate enhancer reporter activity, we mapped and normalized RNA and DNA-derived sequencing reads as counts per million (CPM) for each enhancer-gRNA pairing (summing across barcodes associated with the same gRNA) in each of the 12 experimental samples, and then calculated the RNA-to-DNA ratio. For example, an enhancer fragment from chr1:2187281-2187481 was strongly active in the assay, and the effect was consistent across all 12 samples, with a median activity of 1.68 (log2 (RNA CPM/DNA CPM)) compared to a median activity of −2.24 for scrambled enhancer sequences, *i.e.* 15.1-fold reporter activation (**Fig. 2a**).

**Fig. 2:**
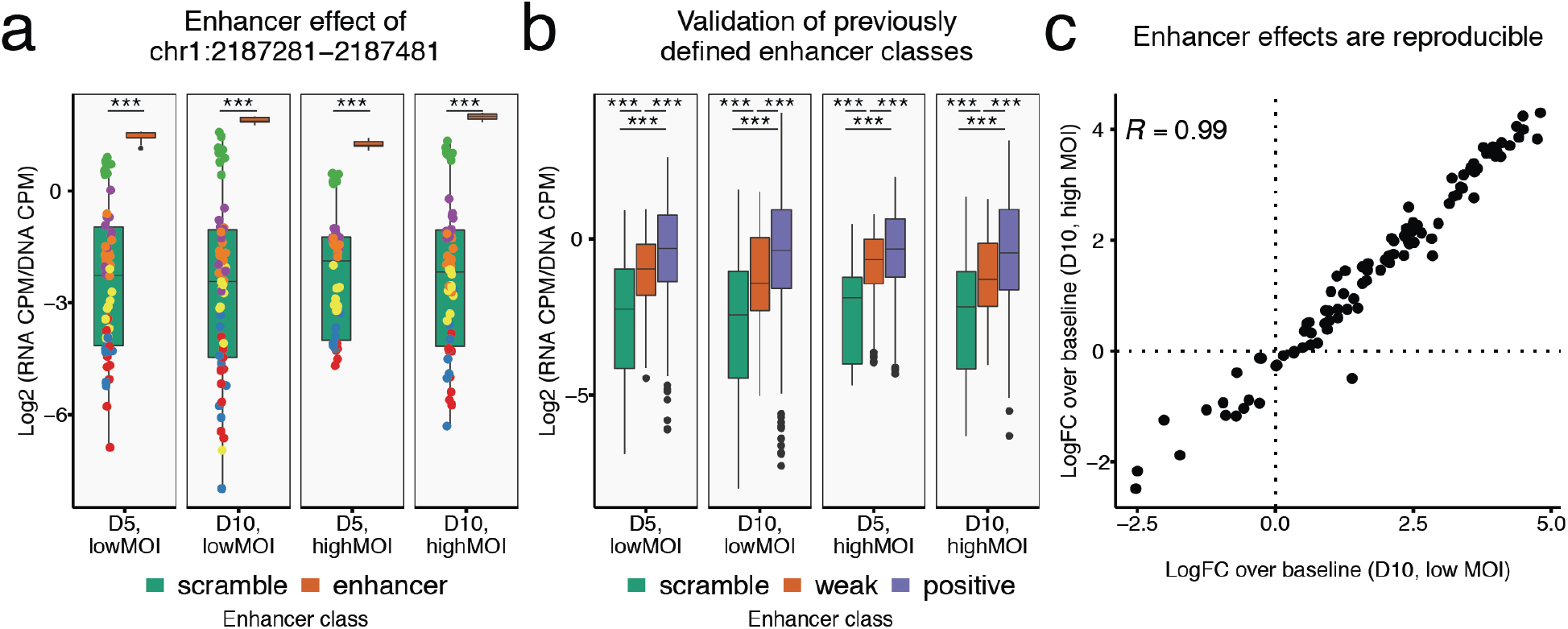
Identifying regulatory regions with piggyBac-mediated *trans*MPRA. **a**, Comparison of reporter activity for scrambled enhancer fragments (green box plot) versus a selected enhancer fragment (chr1:2187281-2187481; orange box plot) for each of the experimental conditions. Colored points on green box plots correspond to individual values for different scrambled enhancers. All pairwise experimental comparisons show this enhancer fragment as having strong activity relative to the scrambled enhancers. Of note, one scrambled enhancer consistently exhibited appreciable activity (green points). *** significant at *P* < 0.001; two-sample T-tests. **b**, Reporter transcription activity for all test DNA fragments grouped by *a priori* assigned enhancer class^24^: scrambled control (green), weak regulators (orange), and positive regulators (blue). *** significant at *P* < 0.001; two-sample, one-sided T-tests. **c**, Reproducibility of enhancer log2-fold-change (“logFC”) over baseline reporter activity (defined as mean activity of scrambled enhancers with scrambled gRNAs) between the high MOI and low MOI conditions sampled from D10. Only enhancers with significant effects above or below baseline reporter activity in either or both conditions were used for Pearson’s R computation (69 of 101; uncorrected *P* < 0.001; two-sample T-test).

To assess whether piggyBac-integrated enhancer fragments were behaving similarly to an episomal assay, we grouped 101 tested enhancer fragments by their *a priori* designation^24^ of ‘scramble’, ‘weak regulator’, or ‘positive regulator’. Across all enhancers paired with scrambled gRNAs, we observed a median 2.24-fold reporter activation relative to ‘scramble’ class enhancers for the ‘weak’ class and median 3.67-fold activation for the ‘positive’ class, relative to the median ‘scramble’ enhancer (**Fig. 2b**). Reassuringly, the results were highly reproducible across conditions, indicating that neither low vs. high MOI nor collection 5 vs. 10 days post-transfection had a major impact on the MPRA itself (**Fig. 2c**; **Fig. S7**). Taken together, we conclude from these analyses that similar to episomal and lentiviral MPRA^27–29^, piggyBac-integrated reporter constructs can successfully and reproducibly identify regulatory enhancers.

We next aimed to identify specific *trans*-acting factors that are relevant to the activity of individual enhancer regions (**Fig. 3**). For this analysis, we compared the activity of specific enhancers paired with scrambled gRNAs vs. the same enhancer paired with a TF-targeting gRNA. For example, we found that the chr1:2187281-2187481 enhancer (**Fig. 2a**) exhibited ∼40% reduced activity when paired with a gRNA encoding CRISPRi of GATA1 (**Fig. 3a**). The observed effect was consistent across all conditions, timepoints and replicates. Notably, while there was no match for the GATA1 motif in this enhancer’s primary sequence, ChIP-seq data supports GATA1 localization to this region in K562 cells (**Fig. S8**).

**Fig. 3:**
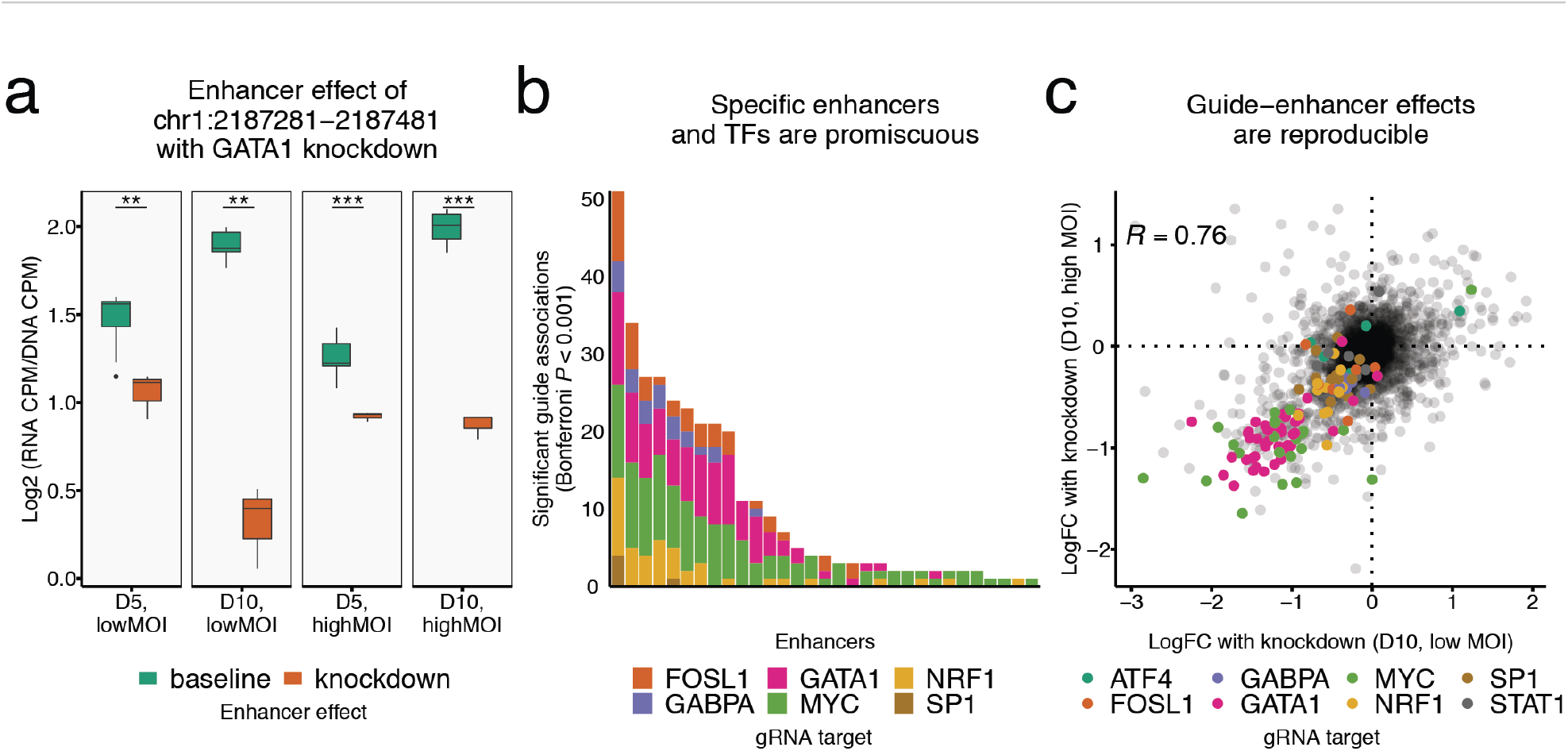
*Trans*MPRA identifies TF knockdown effects on enhancer activity. **a**, Comparison of reporter activity for a selected enhancer fragment (chr1:2187281-2187481) on constructs with scrambled gRNAs (green box plots) versus GATA1-targeting gRNAs (orange boxplots) for each of four sets of conditions. ** significant at *P* < 0.01; *** significant at *P* < 0.001; two-sample T-test. **b**, Distribution of 326 significant guide-enhancer associations across the tested enhancers, out of 31,512 tested interactions (101 enhancers x 26 guides x 2 timepoints x 2 conditions x 3 replicates). We did not observe any significant guide-enhancer associations for 2 of the 8 TFs (ATF4 and STAT1) and 70 of the 101 tested enhancers. **c**, Reproducibility of guide-enhancer knockdown log2-fold-change (“logFC”) effects between high MOI vs. low MOI conditions sampled from D10. Only guide-enhancer combinations with significant effects in either or both conditions were used for Pearson’s R computation (125 of 2,626; uncorrected *P* < 0.001; two-sample T-test).

In total, across 31,512 tested guide-enhancer interactions (101 enhancers x 26 guides x 2 timepoints x 2 conditions x 3 replicates), we identified 329 significant effects (Bonferroni corrected *P* < 0.001). Of the 95 non-scrambled enhancers, 30 had one or more significant interactions with knockdown of one of the eight TFs. Specific enhancers accounted for a disproportionate number of the interactions (**Fig. 3b**). For example, the chr1:2187281-2187481 enhancer exhibited interactions with 6 of the 8 tested TFs. More active enhancers generally had more associations; this is at least partly explained by power, but there were also active enhancers with few or no interactions (**Fig. S9**). Specific TFs accounted for a disproportionate number of interactions. Most notably, guides that targeted MYC or GATA1 for knockdown were associated with significantly reduced activity of 28/95 and 18/95 enhancers, respectively, consistent with their roles as master regulators of gene expression in K562 cells^30^.

The effect sizes of significant guide-enhancer associations were generally reproducible between experimental conditions, particularly between low MOI vs. high MOI experiments, indicating that “cross-reporter” effects within cells with multiple integrants are not substantially impacting the results presented here (**Fig. 3c**). However, D5 estimates were less stable than D10 estimates, which may be due to the time necessary for a given protein-enhancer dynamic to reach equilibrium (**Fig. S10**).

Although we observed an overall strongly significant correlation between the presence of the motif for a given TF in a given enhancer and the detection of a significant interaction, motifs were only weakly predictive (*P* = 9.9 x 10^−7^; one-sided Wilcoxon rank-sum test; **Fig. S11**). For example, there were 10 enhancers with a GATA1 motif match, but we only observed a significant effect for one of these. On the other hand, there were 17 enhancers for which we detected a significant GATA1 interaction despite the absence of a motif.

Using GATA1 as an example, ChIP-seq signals were significantly correlated with interactions, but again only weakly predictive (*P* = 4.1 x 10^−6^; one-sided Wilcoxon rank-sum test; **Fig. S12**). Specifically, there was ChIP-seq evidence for GATA1 binding at the endogenous coordinates of 16 of the 95 enhancers, 5 of which exhibited significant interactions with GATA1 knockdown. However, there were 13 enhancers for which we detected effects despite the absence of ChIP-seq evidence for GATA1 binding. These results suggest a potentially higher-order role for GATA1 (and MYC, which was similarly promiscuous) in enhancer-based gene regulation in K562 cells.

In summary, to enable the quantification of the role of specific *trans*-acting regulatory factors in mediating enhancer effects, we developed the “*trans*” massively parallel reporter assay or *trans*MPRA. As a proof-of-concept, we tested potential interactions between 95 enhancers and knockdown of 8 TFs for effects on reporter transcription. Our results are most analogous to ChIP-seq in that *trans*MPRA has the potential to identify factors with both direct and indirect effects, much as ChIP-seq can detect both direct and indirect binding. However, in contrast with ChIP-seq, *trans*MPRA does not require an antibody and detects functional rather than biochemical effects, including those for which colocalization goes undetected for technical reasons (*e.g.* transient binding) or is biologically unnecessary (*e.g.* protein kinases that modify the activity of TFs). As a functional assay that can be extended to any CRISPR-targetable protein, *trans*MPRA provides an orthogonal avenue for identifying the general and specific *trans*-acting factors underlying gene regulation at *cis* regulatory elements.

From a technical perspective, *trans*MPRA is efficient and flexible. The efficiency arises from linking the measurement of the programmed perturbation and its effect on the same sequencing read^31–33^. In terms of flexibility, one can easily alter the gene-perturbation effect, gRNA targets, enhancer fragments, reporter gene structure, or a variety of other experimental parameters to investigate a broad range of questions about how *trans-*acting factors shape gene regulation.

## Methods

### Identifying putative enhancer regions and selecting TF targets

We downloaded previously collected MPRA data from K562 cells comprising per base reporter activity score for a set of regions assayed through tiling^24^. The regions were subsetted to only include those belonging to the enhancer state (‘5’). To select fragments for the ‘weak regulator’ class, we selected tiled regions that had the lowest max reporter activity score. The putative enhancer was then centered on the base in these tiled regions with the lowest reporter activity score. Flanking regions of length 100 bp were included for a fragment with a total length of 201 bp. To select fragments for the ‘positive regulator’ class, we selected tiled regions with the highest max reporter activity score. The putative enhancer was then centered on the base in these regions with the highest transcription rate score. Again, flanking regions of length 100 bp were included for a fragment with a total length of 201 bp. Finally, to select fragments for the ‘scramble’ class, we took 6 of the previously defined enhancer fragments and randomly permuted the base positions – a process which maintains the proportions of distinct bases while presumably eliminating any enhancer structure. Of the 6 scramble enhancers, 3 were permuted from fragments belonging to the ‘weak regulator’ class and 3 were permuted from fragments belonging to the ‘positive regulator’ class. For Gibson assembly during the iterative cloning process, we included 30 bp of homology sequence on both ends of each putative enhancer fragment for a total length of 261 bp.

We selected TFs to target for CRISPRi knockdown from evidence of PWM-based motif matches within putative enhancers and the expression of TFs in K562 cells. The motif match score or predicted DNA binding affinity of distinct TFs was computed with the ‘motifmatchr’ R package (https://github.com/GreenleafLab/motifmatchr) using default parameter values, which serves as a wrapper to the MOODS motif matching suite. We tested for matches using the ‘human_pwms_v2’ set of PWMs included in the chromVARmotifs package (https://github.com/GreenleafLab/chromVARmotifs). Target TFs were selected based on manual inspection of TF motif matches at putative enhancers. We next verified that the target TFs were expressed in K562 cells and there was evidence of ChIP-seq binding at putative enhancer regions. For both of these analyses, we relied on publicly available ENCODE data visualized with the WashU Epigenome Browser (https://epigenomegateway.wustl.edu/). Once we chose specific TFs to target with CRISPRi, we used an existing library of optimized guides^26^ to select 3 gRNA sequences per target TF. Additionally, we included 3 scrambled gRNA controls that were included in the gRNA library. To simplify the iterative cloning we include several constant fragments to each gRNA. At the end of each fragment we included 30 bp of homology sequence for Gibson assembly. Following the gRNA fragment we included a Cas9 scaffold, a spacer cloning site, and a unique barcode. However, during the cloning process we eliminated the barcode and included random barcodes instead.

The putative enhancers and gRNA fragments were snythesized as two separate oPools at Integrated DNA Technologies (IDT). All fragments described above along with flanking constant regions are listed in **Table S1**.

### Iteratively cloning the paired enhancer-guide transMPRA library

Starting with the piggyBac cargo plasmid (Systems Bioscience PB510B-1), we performed a double digest with SfiI (NEB R0123S) and NheI-HF (NEB R3131S) restriction enzymes. A custom gBlock (**Table S1**) with a U6 Pol3 promoter, a cloning site containing two BseRI cut sites, and a SV40 polyA signal were cloned into the digested plasmid with NEBuilder HiFi DNA Assembly (NEB E2621). The resulting product was transformed into stable chemically competent *E.coli* (NEB C3040H) and plated. Several individual colonies were isolated, grown, maxi-prepped (Zymo D4202), and verified with Sanger sequencing.

We digested the resulting plasmid with BseRI (NEB R0581L) and agarose gel-size selected the linearized fragment. The custom DNA fragment with the gRNA library, a Cas9 scaffold, spacer cloning site with two BseRI cut sites, and custom designed barcode was amplified with library-specific primers and the KAPA HiFi HotStart ReadyMix (Kapa KK2602) and then was agarose gel size-selected. The size-selected fragment was cloned into the digested plasmid with NEBuilder HiFi DNA Assembly (NEB E2621), and the resulting product was transformed into 10-Beta Electrocompetent cells (NEB C3020K). We plated 1% of the library to estimate complexity and grew the rest of the sample and then midi prepped (Zymo D4200) the resulting library.

The gRNA library was amplified with a primer that included an NheI cut site. The amplified library was then cloned into the previously digested plasmid, and then the resulting library was midi prepped. To add a random barcode, we digested this plasmid with NheI-HF and then cloned a custom DNA primer with an 18 bp random barcode with NEBuilder HiFi DNA Assembly. The library was then transformed into 10-Beta Electrocompetent cells and midi-prepped. Further below, we describe our sequencing strategy for associating random barcodes with guides.

Following the inclusion of the random barcodes, we digested the plasmid library with BseRI (NEB R0581L) and agarose gel size-selected the linearized fragment. The ORI minimal promoter and flanking region was PCR amplified from the hSTARR-seq plasmid (Addgene #99296) with a custom primer that included homology for Gibson cloning and KAPA HiFi HotStart ReadyMix (Kapa KK2602) and then was agarose gel size-selected. We then included the minimal ORI promoter and flanking region between the gRNA and random barcode with NEBuilder HiFi DNA Assembly. Again, the plasmid library was then transformed into electrocompetent cells and then midi prepped.

For the final step, the previous plasmid library was digested with BseRI and the linearized fragment was agarose gel size-selected. The custom DNA fragment pool with putative enhancers was amplified with library-specific primers and KAPA HiFi HotStart ReadyMix, and then agarose gel size-selected. We used NEBuilder HiFi DNA Assembly to include the enhancer library into the BseRI-digested vector that already included the gRNA, random gRNA-linked barcode, and minimal ORI promoter. Again, the plasmid library was then transformed into electrocompetent cells and then midi prepped.

### Associating guides and barcodes through deep sequencing

At the cloning step before the incorporation of the minimal promoter, we deeply sequenced the plasmid library to associate guides with random barcodes. From this plasmid library, we PCR-amplified the section of interest with two amplicon-specific primers that incorporate a specific adapter sequence. We performed a subsequent PCR amplification to add sample indices and the P5 and P7 flow cell adapters. Products were pooled with other samples on a NextSeq instrument. This library was sequenced twice to increase the number of barcode-guide associations.

Overall, we collected 40 million reads that passed QC. Reads were aligned with bowtie2 version 2.3.5. In preparation for alignment, two bowtie indices were built with default parameters – one index based on the amplicon sequence where the barcode positions were replaced with ‘N’s and another index based on the amplicon sequence with one version per guide. The read fragment fastq files including the barcode segment were aligned to the barcode-specific bowtie index with ‘--n-ceil L,18,0.15’ and otherwise default parameters. The read fragment fastq files containing the gRNA sequence were aligned to the gRNA-specific bowtie index with default parameters. From the bam output of these alignments, for each read we extracted the gRNA fragment which the read aligned to and the random barcode sequence. We excluded read fragments with Ns in the barcode and fragments that had barcodes paired with multiple guides. In total, we identified ∼1.75 million unique pairs of barcodes and guides.

### Cell culture and transformation

K562 cells are derived from a female with chronic myelogenous leukemia and are an ENCODE Tier 1 cell line. The Bassik lab gifted us K562 cells that were transduced to express dCas9-BFP-KRAB (Addgene #46911, polyclonal). The cells were grown at 37°C and cultured in RPMI 1640 with L-Glutamine (GIBCO) along with 10% FBS and 1% penicillin-streptomycin (GIBCO). Cells were confirmed to express BFP with FACS.

To transduce custom DNA fragments into cells we used the piggyBac transposase system, which relies on co-transfecting the DNA library cloned into the transposon cassette (the product of our iterative cloning process) along with the piggyBac transposase vector (Systems Bioscience PB210PA-1). Our approach requires low integration rates per cell so as to avoid inhibiting cell proliferation and avoid the prevalence of cells with many plasmids that target distinct TFs. Therefore, we first set out to optimize the ratio of transposase to transposon that correspond with specific rates of integration. For this optimization experiment we co-transfected the transposase with a GFP gene included in the piggyBac transposon cassette. The GFP plasmid and transposase were co-transfected with the MaxCyte STX electroporation system (MaxCyte Systems) as per the manufacturer’s guidelines. **Table S1** lists the distinct transfection conditions tested. Transformed cells were passaged normally and aliquots were taken at day 2, 6, 8, and 10 post transfection for FACS analysis using a FACSAria II (Becton Dickinson).

We determined the proportion of GFP-expressing cells for samples with the different transfection conditions, including a control which excluded the transposase (**Fig. S3**). Assuming a transposase integration follows a Poisson process we can back calculate the average number of integrations per cell (referred to as MOI) following existing approaches^34^. From these data we decided to experimentally test two conditions with our *trans*MPRA library: 1) a ‘highMOI’ condition with 5 ug of *trans*MPRA library and 30 ug transposase (with an estimated MOI of ∼2); and, 2) a ‘lowMOI’ condition with 1 ug of *trans*MPRA library and 30 ug of transposase with an unknown MOI, but likely lower than 2 since it represents 20% of the amount of library for the ‘highMOI’ condition. Additionally, we chose to examine samples from days 5 and 10 post transfection.

Using the lowMOI and highMOI condition defined using the GFP optimization experiment, we co-transfected the *trans*MPRA library along with piggyBac transposase with conditions described in **Table S1** as per the manufacturer’s guidelines. Following transfection the cells were passaged normally. Aliquots of 5 million cells were harvested on day 5 and 10 post transfection and immediately processed upon harvest.

### Cell sample processing and sequencing

Following our experimental design (**Fig. S4**; **Table S1**) at day 5 and day 10 post transfection, cells were harvested, genomic DNA and total RNA were extracted using the AllPrep DNA/RNA mini kit (Qiagen 80204). We extracted mRNA from total RNA with the Oligotex Direct mRNA mini kit (Qiagen 72022).

We used a One-Step RT-PCR kit (Thermofisher 12595025) with custom primers to produce cDNA from the mRNA and subsequently ran 3 cycles of PCR which included a P5 adapter, sample-specific p5 index (8 bp), UMI (10bp) and P7 adapter (**Fig. S5**). RT-PCR products were cleaned with AMPure XP beads (Beckman Coulter A63880). Next, the library was amplified using P5/P7 primers. Finally, the resulting PCR-amplified cDNA library was pooled at an equimolar ratio then agarose gel size-selected.

DNA was processed with a similar two-step PCR approach. First, we PCR amplified the DNA for 3 cycles, which incorporated a P5 adapter, sample-specific p5 index (8 bp), UMI (10 bp), and P7 adapter. PCR products were cleaned with AMPure XP beads (Beckman Coulter A63880). Next, we amplified the library using P5/P7 primers. Finally, the resulting PCR-amplified DNA library was pooled at an equimolar ratio and then agarose gel size-selected.

All RNA and DNA libraries were pooled and sequenced on an Illumina NextSeq instrument. Paired-end reads of 150 base pairs were sequenced from the forward and reverse end of the amplified fragment. Reads from specific enhancer-barcode plasmids were collapsed by UMI to avoid PCR amplification biases.

### Read alignment and count processing

Before aligning reads to the construct, overlapping reads were merged with pear^35^ version 0.9.10 with the flags ‘-n 240 -m 260’ and otherwise default parameters. For alignment we constructed a bowtie2 (version 2.2.5) index with default parameters using a fasta file generated from the construct sequence with a distinct sequence for each enhancer and ‘N’ values at the random barcode region. The bowtie2 alignment was performed with ‘--threads 4’, ‘--n-ceil L,18,0.15’, and the pear-merged reads as the input. Following read alignment we summarized each read by the enhancer it best aligned to as well as the random barcode sequence. We used the file of unique guide and barcode pairs described above to perfectly match barcodes to guides.

At this point we saved two sets of summary data. We saved all the count values for all samples without aggregating by barcodes that uniquely identify the guide (**Supplementary Data 1**). Additionally, we created a count matrix for the samples where we summed the number of counts that associate with a guide and enhancer pair, essentially summing across barcodes (**Supplementary Data 2**). Both summary data sets were normalized to account for read depth in the same way. We primarily visualized the summary data aggregated across barcodes but used the full barcode data to perform hypothesis testing for each sample (described in further detail below).

After collapsing by UMIs, we used the calcNormFactors function with default parameters and the cpm function from edgeR version 3.26.8 to compute for each DNA and RNA sample the number of reads per million aligned fragments (CPM) for each construct with a gRNA. The cpm function by default includes a pseudocount of 2 to handle 0 values.

Genes that affect cell proliferation could adversely affect estimates of transcription if we only measured RNA, for this reason we normalize the RNA CPMs by the DNA CPMs i.e., log2(RNA CPM / DNA CPM). This value represents the normalized reporter activity, which accounts for sequencing depth, differential abundance of plasmids, and proliferation effects.

### Testing for significant transcription rate effects

To test for enhancer effects, we aimed to compare the estimated reporter activity for constructs with a particular enhancer to the baseline reporter activity for the construct with a scrambled control enhancer. For this test, we excluded all constructs without a scramble guide. To identify significant enhancer effects even from a single replicate we considered constructs with distinct barcodes as independent replicates. In parallel, we used the aggregated counts that were computed by summing across barcodes to test for a consistent effect between the three independent replicates. For both cases, we tested for a differential mean transcription rate using a standard T-test implemented in R with the t.test function. Additionally, we included the results from using a nonparametric Wilcoxon rank sum test which correlated with the results from the T-test.

To test for guide-enhancer effects, we aimed to compare the estimated reporter activity for constructs with a particular gRNA and enhancer to the baseline reporter activity for that particular enhancer with scrambled guides. To identify significant guide-enhancer effects even from a single replicate we once again considered constructs with distinct barcodes as independent replicates. In parallel, we used the aggregated counts that were computed by summing across barcodes to test for a consistent effect between the three independent replicates. For both cases, we once again tested for a differential mean transcription rate using a standard T-test implemented in R with the t.test function. Additionally, we also included results from using a nonparametric Wilcoxon rank sum test which correlated with the results from the T-test.

In addition to a p-value, we performed multiple hypothesis test correction with both the Bonferroni and the Benjamini-Hochberg methods (as implemented with the p.adjust function in R) and included these values in the summary data. The p-values included in the figures were uncorrected (unless otherwise stated in the figure legends) as they were computed from examples of tests that were significant following Bonferroni correction when performed on unaggregated counts.

### Publicly available data

Two replicates of ChIP-seq targeting GATA1 in K562 cells were downloaded from the ENCODE data portal in the form of p-values of read enrichment over control samples^25^. We consider GATA1 bound to an enhancer if there was at least one base with *P* < 1 x 10^−5^ ChIP-seq enrichment in both replicates.

## ENDNOTES

### Data availability

The data generated can be downloaded in raw and processed forms from the National Center for Biotechnology Information’s Gene Expression Omnibus (GSE157430). We included normalized reporter activity values (log2(RNA CPM/DNA CPM)) for the unaggregated (**Supplementary Data 1**) and aggregated versions of the data (**Supplementary Data 2**).

## Acknowledgements

We thank Jacob W. Freimer and Silvia Domcke for early discussions, and Koshlan Mayer-Blackwell for preliminary feedback. We are grateful to Michael Bassik’s lab for providing the dCas9-BFP-KRAB-expressing K562 cell line. This work was supported by the National Human Genome Research Institute grants 1UM1HG009408 (J.S.), 5R01HG009136 (J.S.), and 1R01HG010632 (J.S. and C.T.). D.C. and the project described was further supported by Award Number T32HL007828 from the National Heart, Lung, and Blood Institute. J.S. is an Investigator of the Howard Hughes Medical Institute.

## Author contributions

D.C. conceived of the initial idea. D.C., A.E., C.T., and J.S. conceptualized the presented study. C.T. and J.S. supervised the study. D.C., R.M.D., B.M., J.M.T., W.C., F.M.C., and A.L. designed the cloning strategy, tissue culture methods and transfection approach. D.C., R.M.D., B.M., W.C., and C.L. planned and implemented the sequencing strategy. D.C. carried out the data collection, performed the formal analysis, and wrote the original draft. D.C., C.T., and J.S. reviewed and edited the draft. All authors read, provided feedback, and approved the final manuscript.

## Competing interests

The authors declare no competing interests.

## Supplementary Figures

**Fig. S1:**
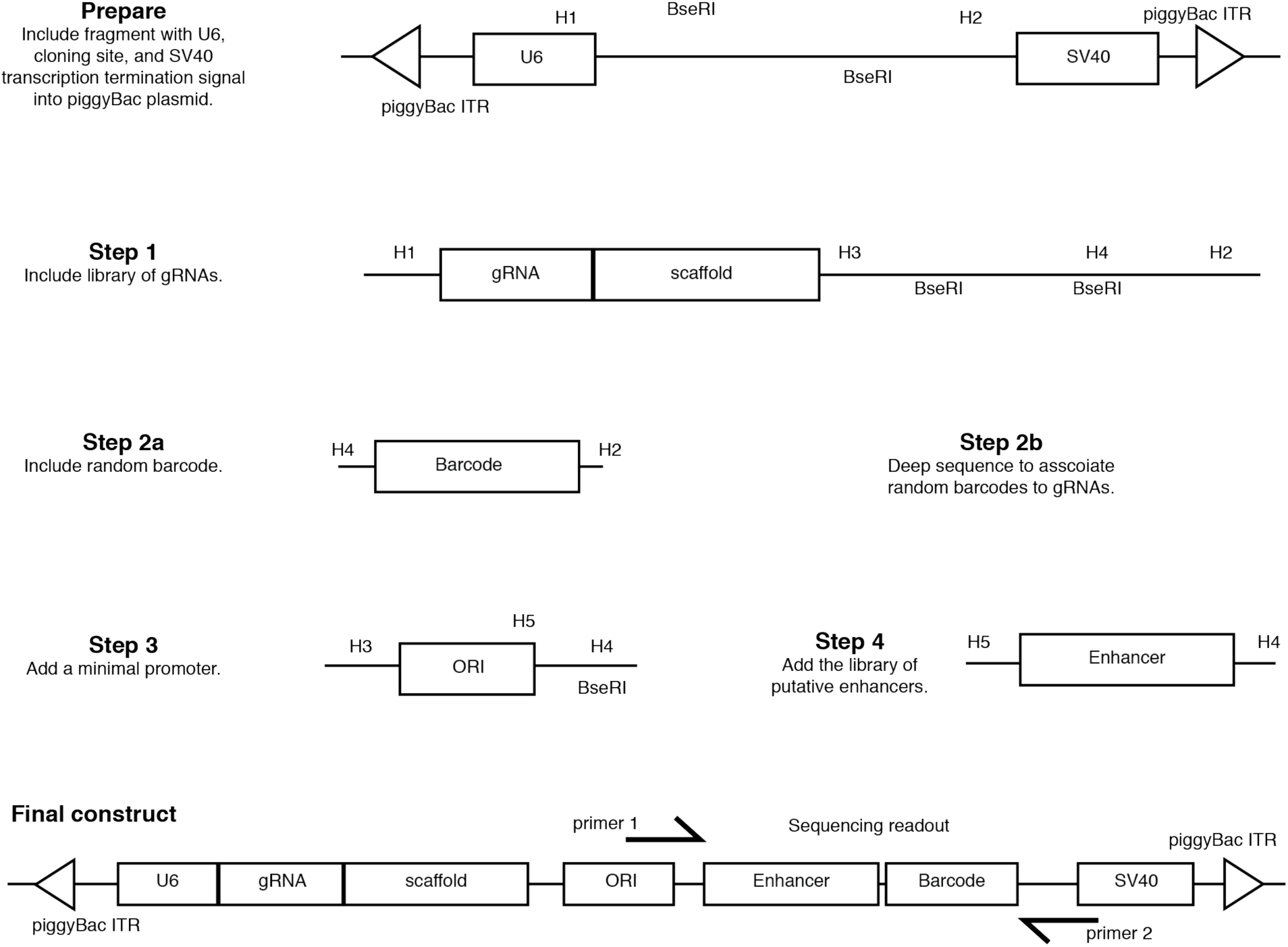
Cloning strategy. First a Pol3-associated U6 promoter and SV40 transcription termination element are cloned into the piggyBac cassette plasmid (“Prepare”). The cloned fragment contains a cloning site with two BseRI cut sites and homology for Gibson assembly. Then a gRNA library along with a scaffold region are cloned into the cloning site (“Step 1”). This fragment also contains a cloning site. A random barcode is added (“Step 2a”). Before continuing, we sequence the amplicon to associate barcodes to guides (“Step 2b”). The minimal promoter is added between the barcode and the gRNA scaffold (“Step 3”). Finally, we clone the library of putative enhancer elements adjacent to the random barcode resulting in the final construct (“Step 4”). Regions labeled with H represent regions of homology used for Gibson assembly.

**Fig. S2:**
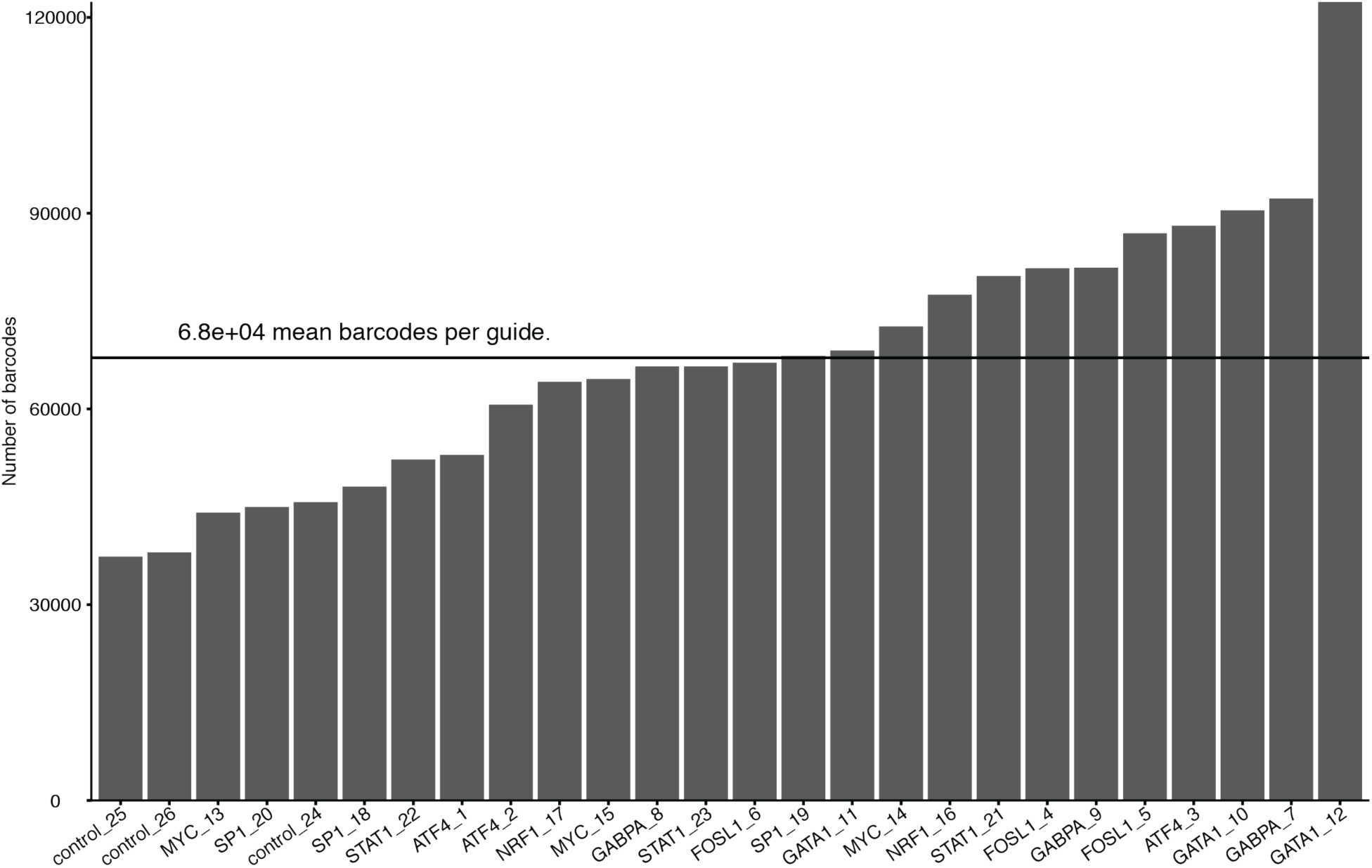
Barcode-guide associations. The number of unique barcodes associated with each of 26 gRNAs. The horizontal line indicates the mean number of barcodes per guide.

**Fig. S3:**
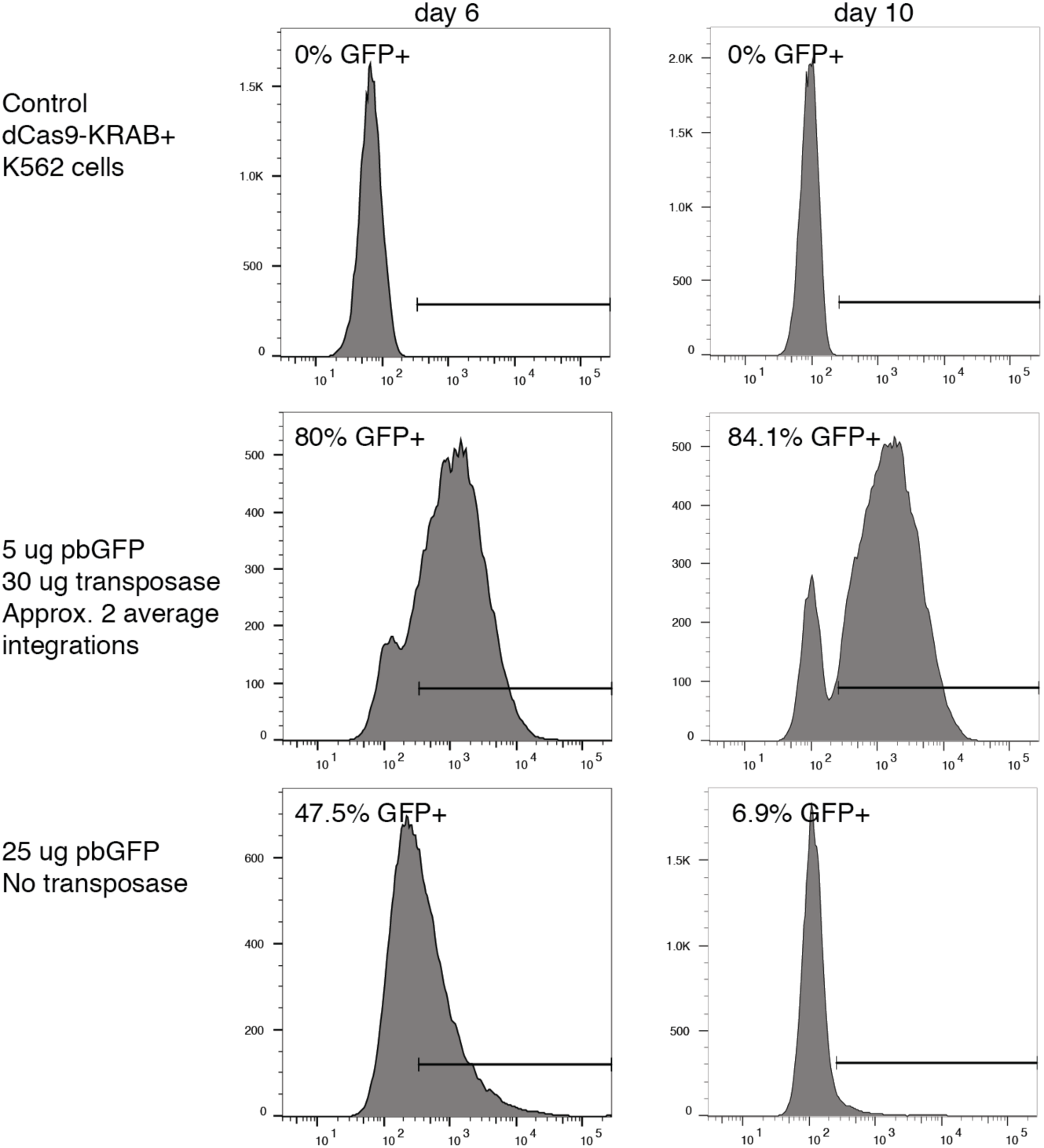
PiggyBac GFP optimization. Distribution of GFP expression among cells transfected at different library concentrations and harvested at two distinct timepoints (day 6 vs day 10) that most closely replicate the chosen experimental conditions. Cells were transfected with a piggyBac transposon containing the GFP gene along with the piggyBac transposase plasmid (except for the bottom control condition). The proportion of GFP expressing cells is indicated.

**Fig. S4:**
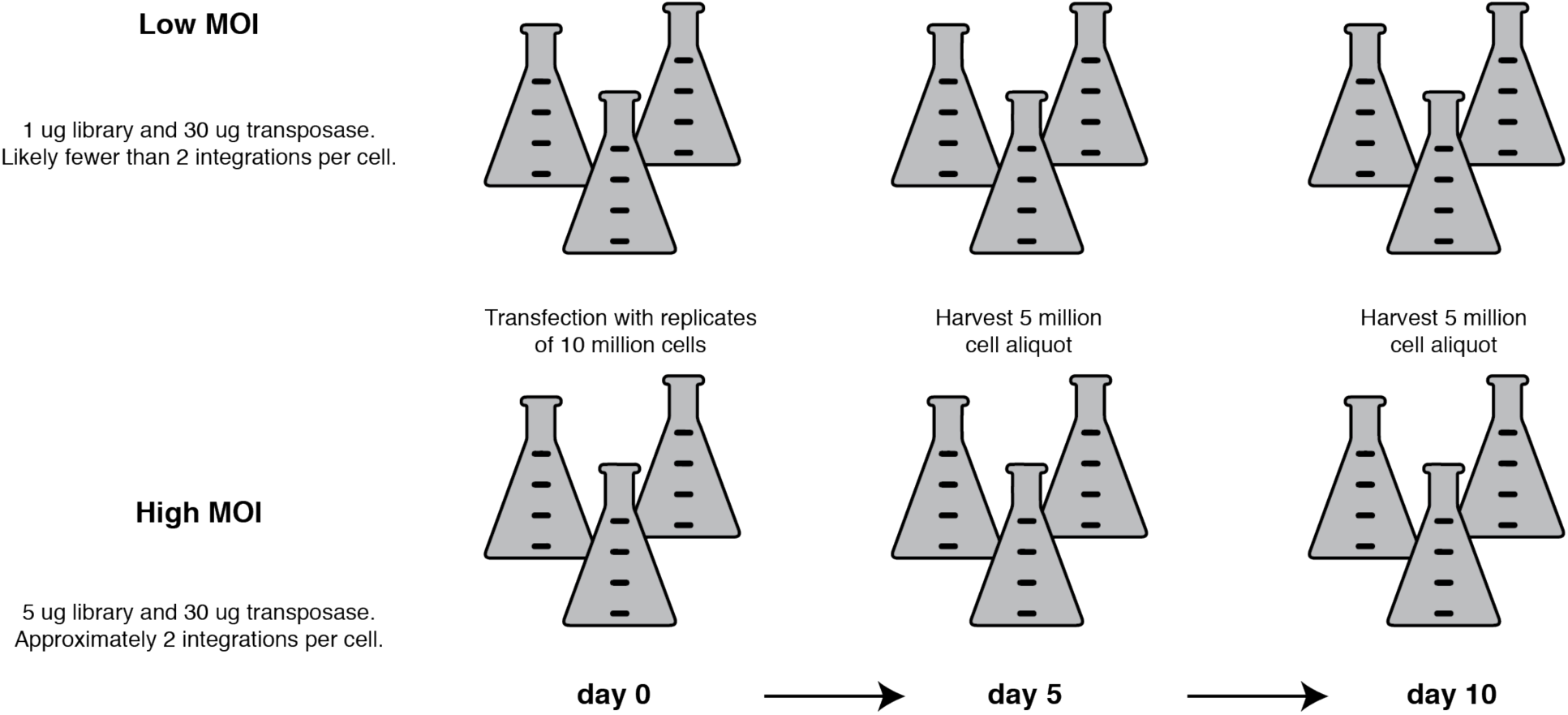
Experimental design. The *trans*MPRA library was transduced into three replicates of 10 million K562 cells engineered to constitutively express the dCas9-KRAB repressive complex. We used 2 library concentrations: a high multiplicity of integration (highMOI) condition and a low multiplicity of integration (lowMOI) condition. Aliquots of 5 million cells were harvested on day 5 and day 10 post transfection.

**Fig. S5:**
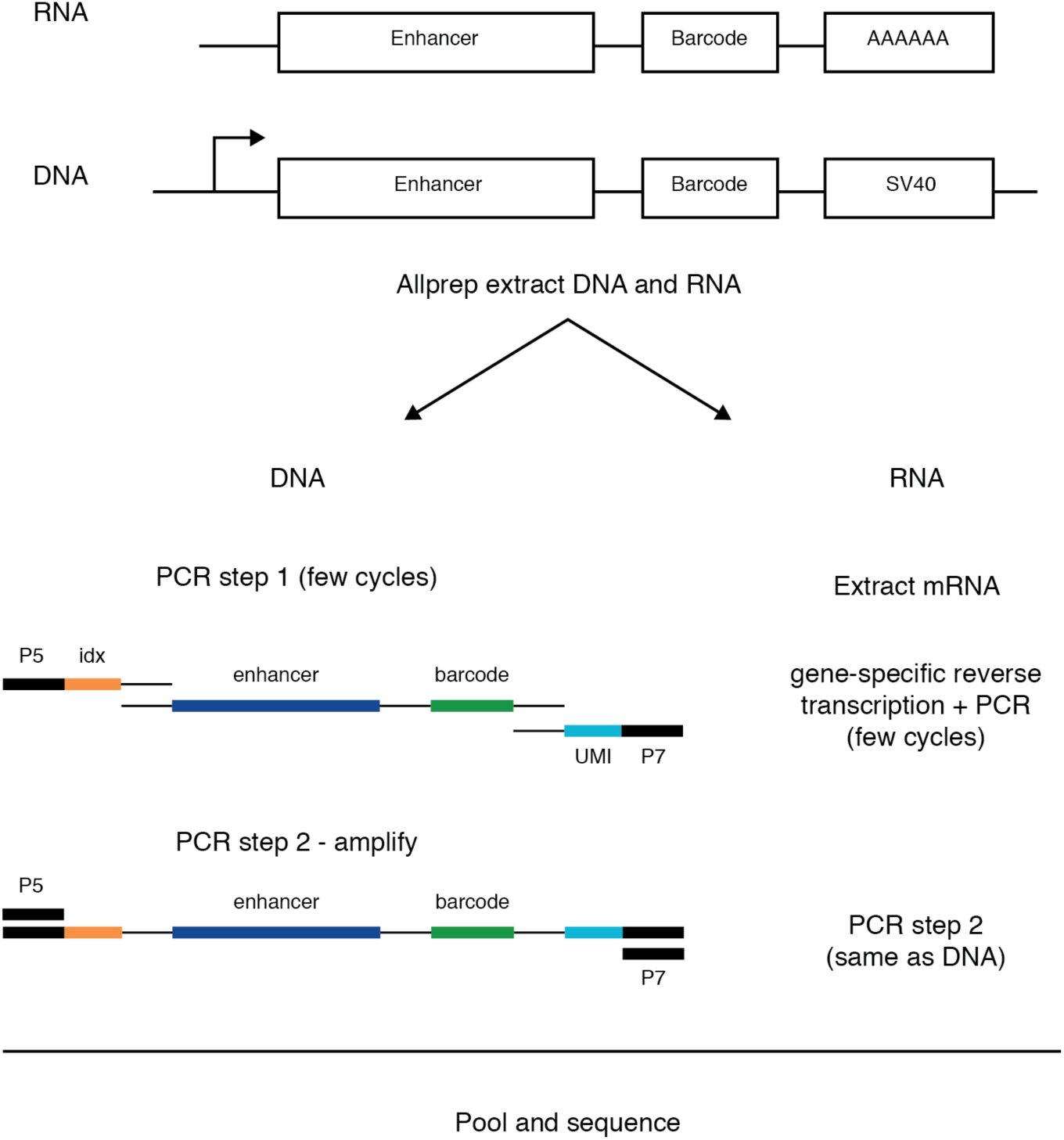
Sequencing strategy. From aliquots of 5 million cells we first extract both DNA and RNA from the cells. We use a two-step PCR strategy to add all relevant indices and adapters. For the DNA, with an amplicon-specific sequence primer, the first PCR adds a P5 flow cell adapter, P5 index, UMI, and P7 flow cell adapter. The second PCR amplifies the fragment using the P5/P7 flow cell adapters as primers. For the RNA we first extract mRNA from the total RNA. Then using the same amplicon-specific primer we perform a one-step RT-PCR that again includes a P5 flow cell adapter, P5 index, UMI, and P7 flow cell adapter. The second PCR step again amplifies the fragment with P5/P7 flow cell primers. The RNA and DNA samples are then pooled by assay type, gel-size selected, and sequenced.

**Fig. S6:**
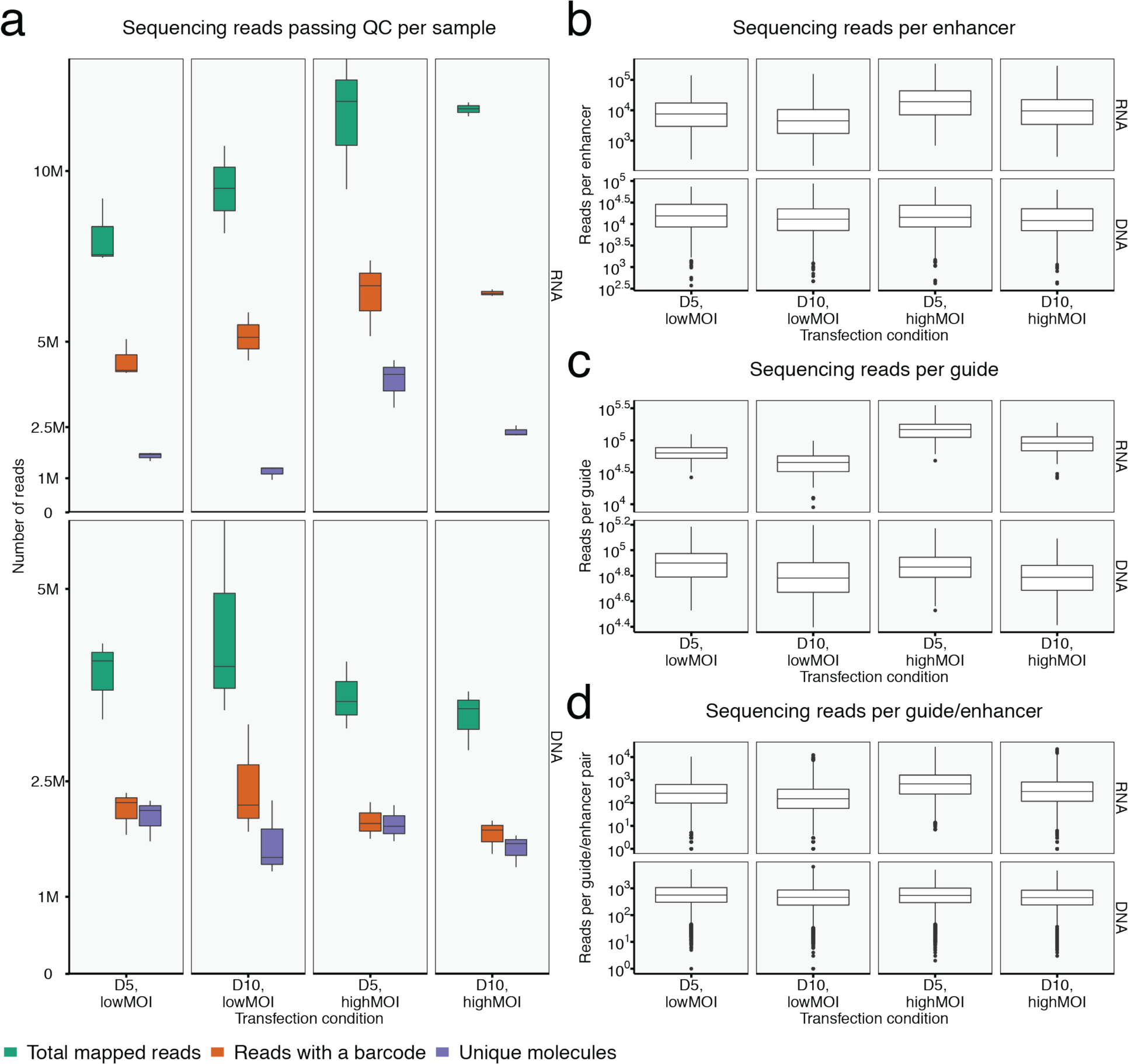
Distribution of read sequencing. **a**, Number of sequencing reads that passed basic QC for each sample. **b**, Number of reads per enhancer for each sample. **c**, Number of reads per guide for each sample. **d**, Number of reads per enhancer guide pair for each sample.

**Fig. S7:**
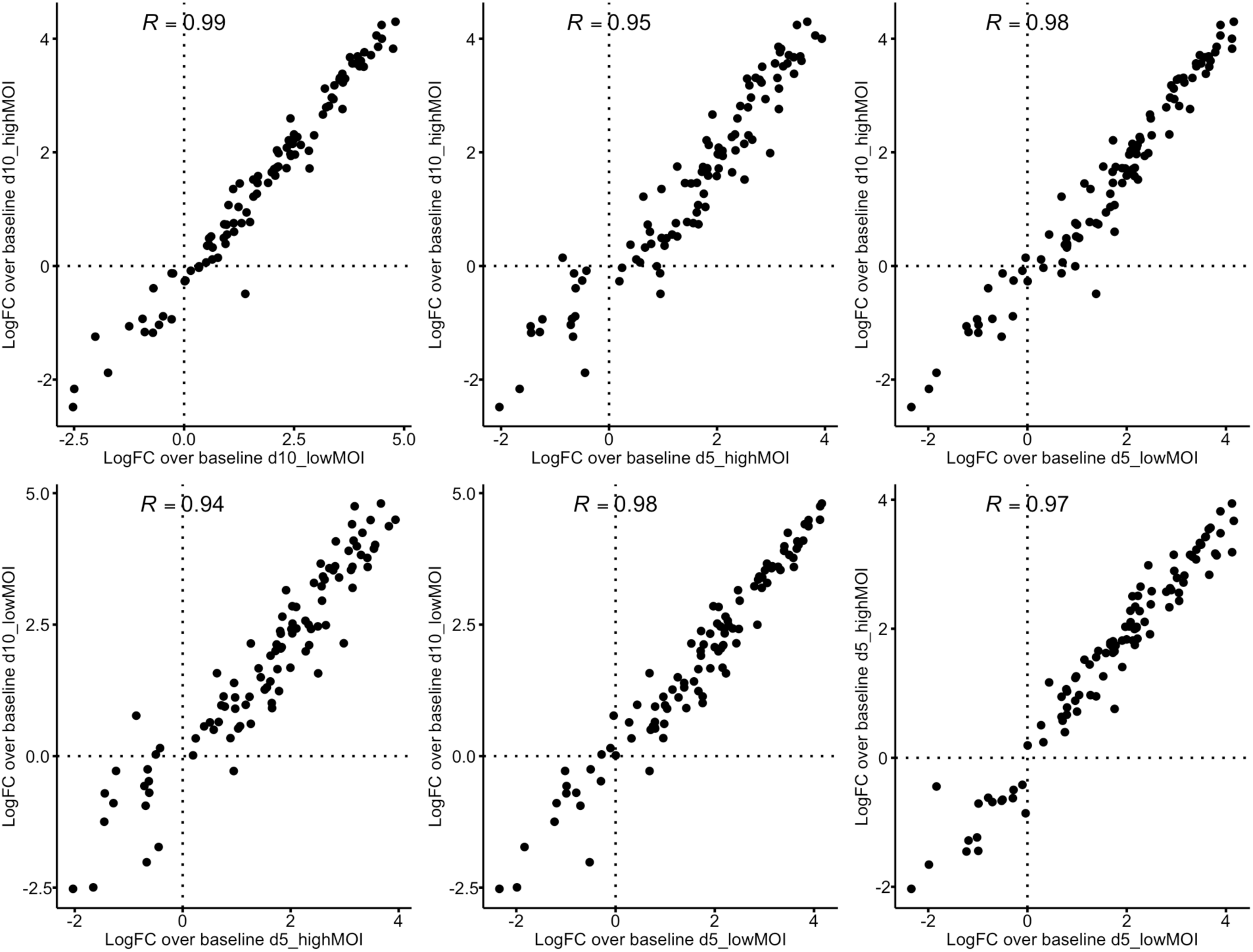
Correlation of enhancer logFC above baseline transcription from transposase-based MPRA analysis. Reproducibility between enhancer logFC above baseline reporter activity from different library concentrations and cells harvested at different time points. Pearsons’s R correlation values were computed from tests marginally significant (*P* < 0.001; two-sample T-test) in at least one of the two conditions compared.

**Fig. S8:**
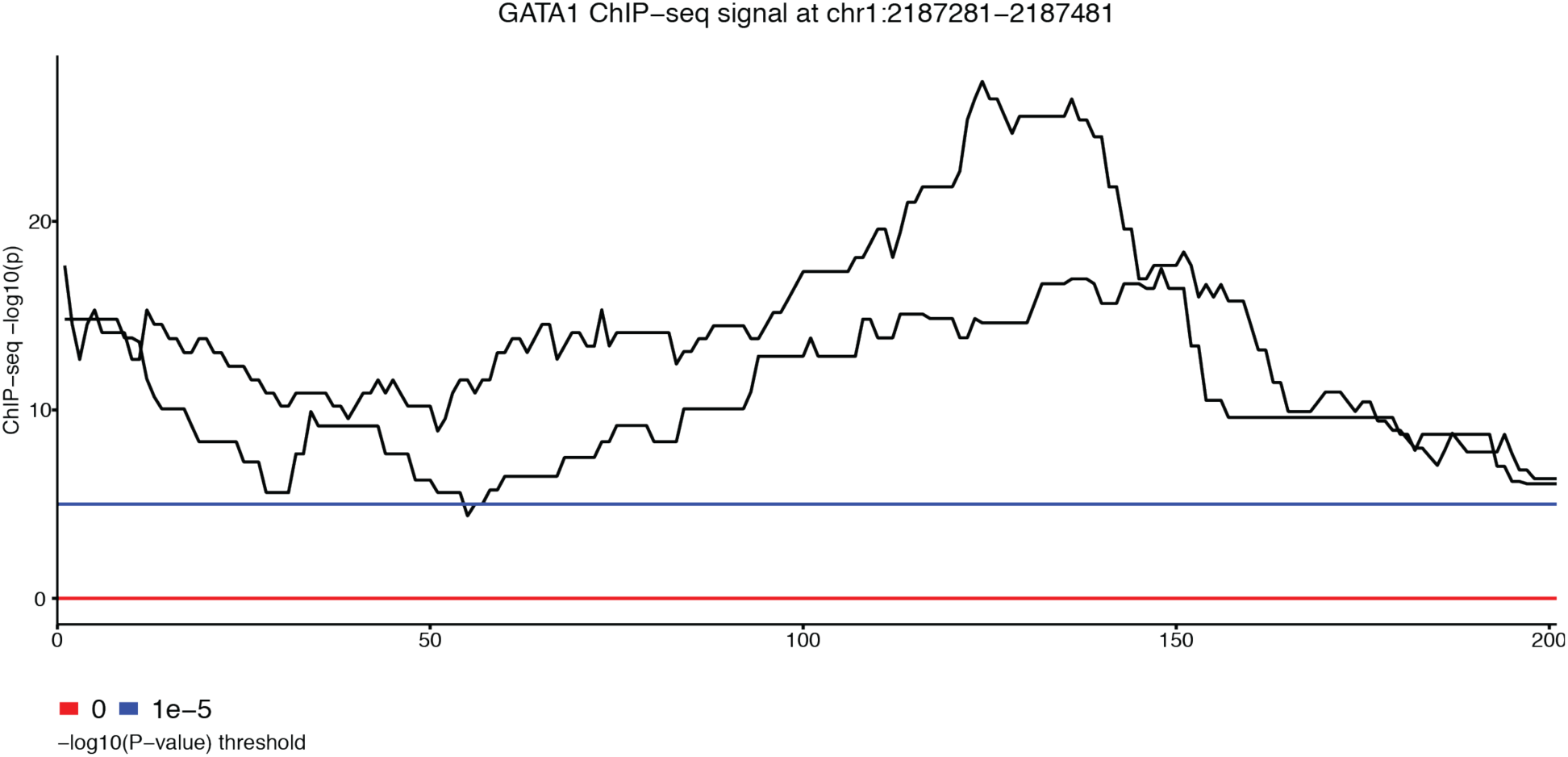
GATA1 binding at an enhancer from chr1:2187281-2187481. Visualization of per base -log(p) enrichment over background of ChIP-seq reads that target GATA1 in K562 cells from 2 replicates across a putative enhancer fragment from chr1:2187281-2187481. A significance threshold of *P* = 1 x 10^−5^ is indicated as a blue line whereas a threshold of *P* = 1 is indicated as a red line. Data were publicly available through the ENCODE data portal.

**Fig. S9:**
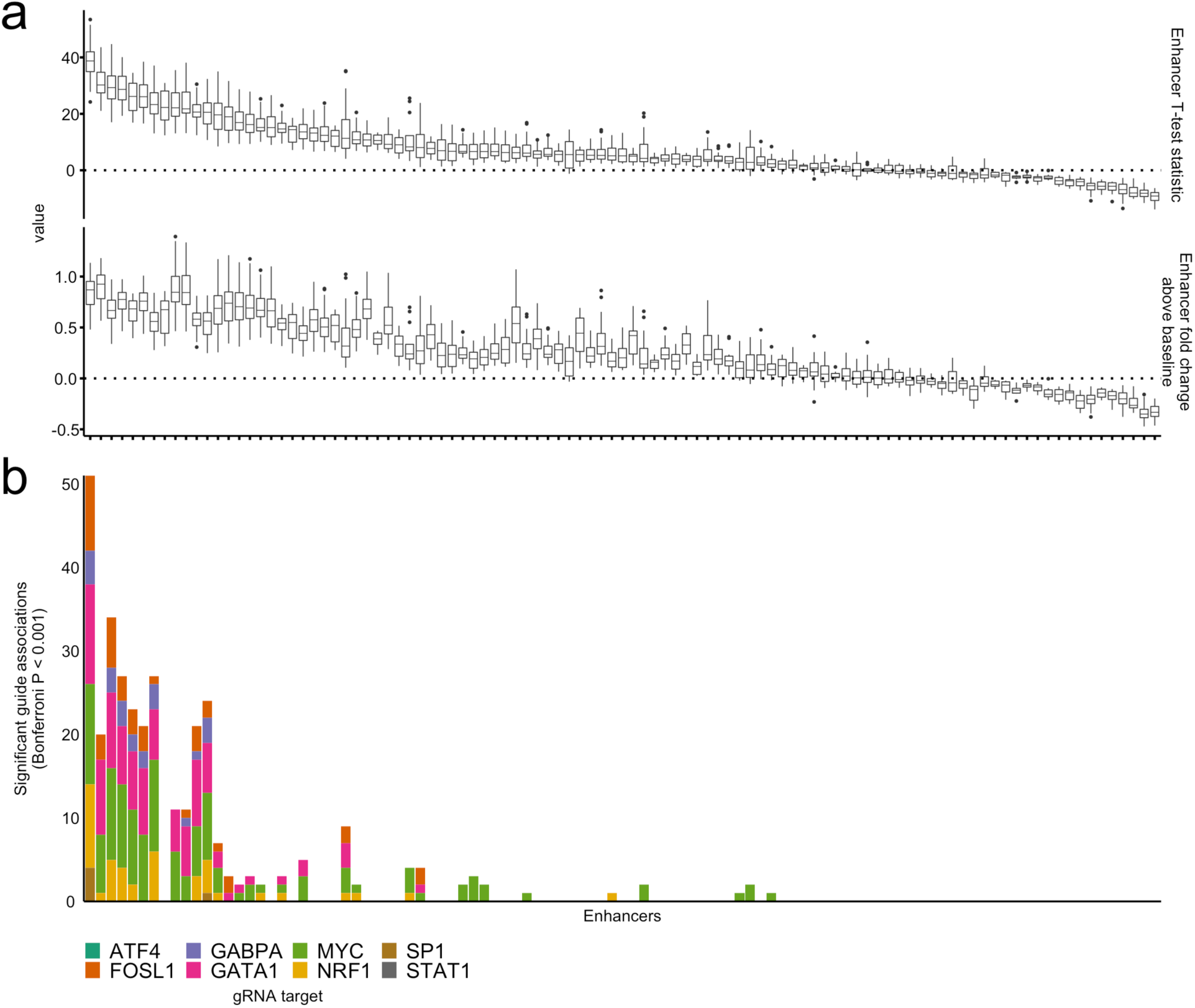
Stronger enhancers have more significant guide associations. **a**, Distribution across replicates of T-test statistics for enhancer effects relative to baseline transcription (top). Distribution across replicates of fold change for enhancers relative to baseline transcription (bottom). **b**, Distribution of the count of significant guide associations per enhancer, which includes enhancers with no observed interactions. Enhancers from the two panels are in the same order. Enhancers are ordered by the median T-test statistic of enhancer-associated reporter activity across replicates.

**Fig. S10:**
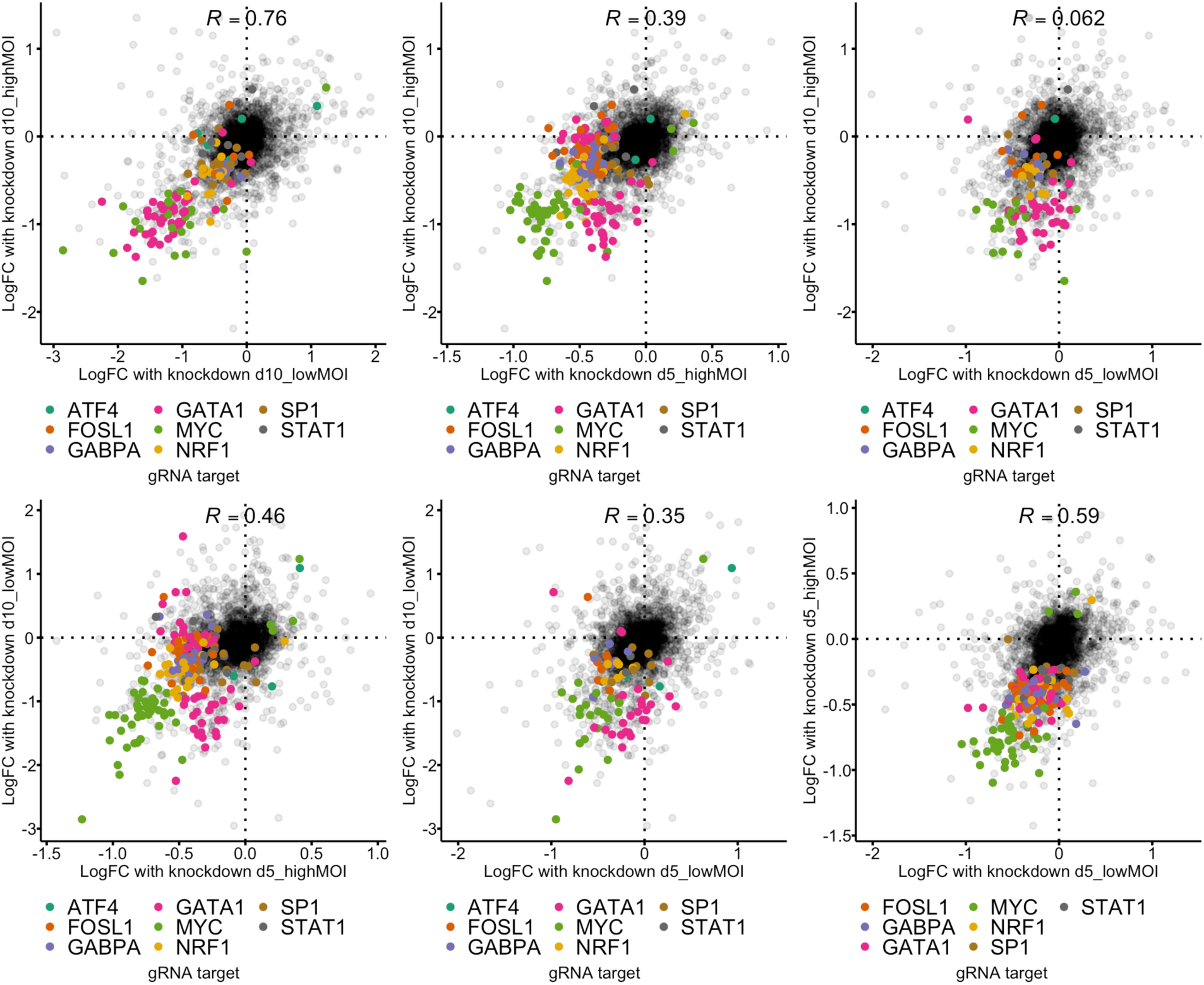
Correlation of effect estimates from guide-enhancer interaction MPRA analysis. Reproducibility of enhancer-guide logFC effects between different library concentrations and cells harvested from different time points. Pearsons’s R correlation values were computed from tests marginally significant (*P* < 0.001; two-sample T-test) in at least one of the two conditions compared. These tests are represented as colored points corresponding with the gene knockdown target.

**Fig. S11:**
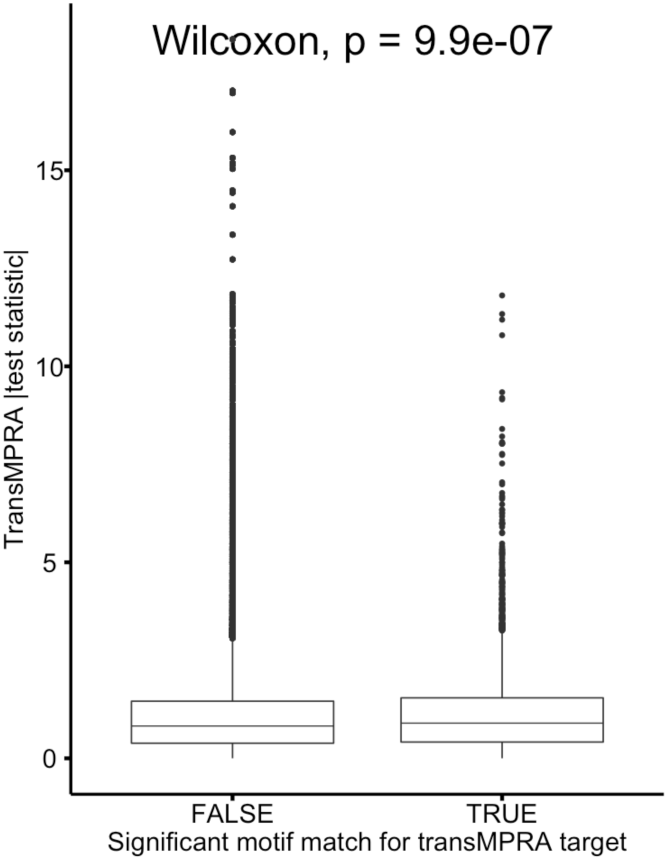
Correlation between motif matches and *trans*MPRA-effect associations. Distribution of absolute value *trans*MPRA T-test statistic across all tests stratified by whether there is a significant motif match for the target gene.

**Fig. S12:**
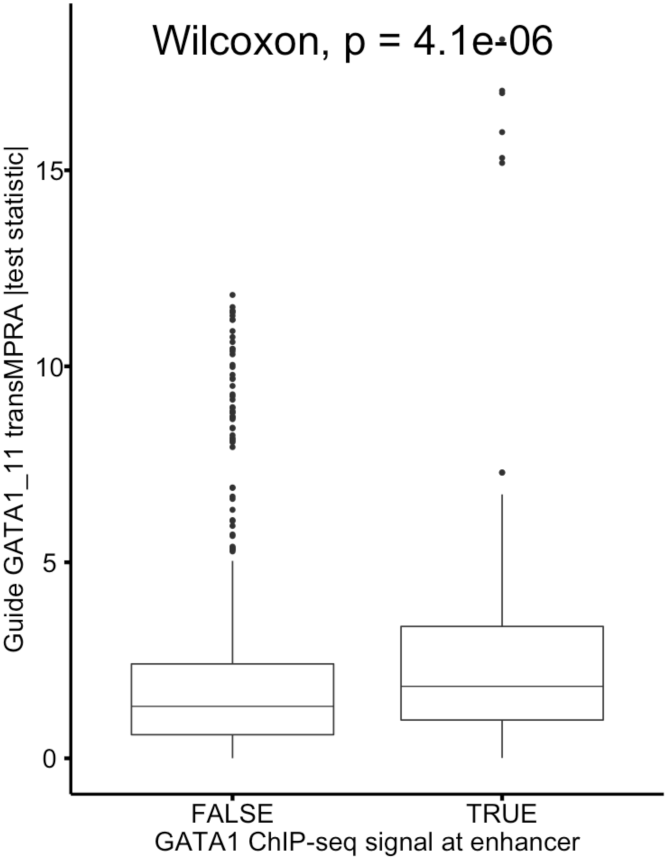
Correlation between GATA1 ChIP-seq and guide-GATA1_11 *trans*MPRA-effect associations. Distribution of absolute value *trans*MPRA T-test statistic across all tests stratified by whether there is evidence of GATA1 binding from ChIP-seq.

## Notes

https://www.ncbi.nlm.nih.gov/geo/query/acc.cgi?acc=GSE157430

### Competing Interest Statement

The authors have declared no competing interest.

